# Virophage-like viruses replicate with a candidate phycodnavirus of *C. parva*

**DOI:** 10.64898/2026.09.29.755237

**Authors:** G. R. Thomas, I. Inamoto, S. M. Short

## Abstract

The discovery of virophages and polinton-like viruses (PLVs) sparked studies of their hyperparasitic relationships with their associated giant viruses and hosts. To date, every cultured virophage and PLV depends on an *Imitervirales* helper virus. The freshwater alga *Chrysochromulina parva* is infected by viruses in the *Imitervirales* and *Algavirales*, and recent work showed that the PLV, CpV-PLV Moe, replicates via co-infection with the *Imitervirales* host CpV-BQ3. The other known PLVs, CpV-PLV Larry and CpV-PLV Curly, were subsequently isolated alongside CpV-BQ1, a putative *Algavirales* virus, raising the possibility that they depend on an *Algavirales* helper. To determine the nature of Larry and Curly’s hyperparasitic relationships, infection experiments were conducted with each PLV and either CpV-BQ3 or CpV-BQ1. These experiments demonstrated that that neither Larry nor Curly replicated when introduced to *C. parva* alone, nor during mixed infections that included the *Imitervirales* virus BQ3. In contrast, the *Algavirales* virus CpV-BQ1 supported replication of both PLVs, with each increasing roughly 1000-fold over the course of infection while CpV-BQ1’s replication was simultaneously reduced up to 100-fold. Together, these findings provide the first culture-based evidence that PLV replication can be supported by an *Algavirales* helper and demonstrate that *C. parva* supports at least two distinct hyperparasitic relationships spanning different viral orders within the class *Megaviricetes*.

**Author summary:** This study revealed that two polinton-like viruses (PLVs), CpV-PLV Larry and CpV-PLV Curly, rely on a previously unrecognized type of helper virus to replicate. PLVs are hyperparasites related to virophages; they infect giant viruses that themselves infect microbial hosts. Until now, all PLVs studied in the lab depended on helper viruses from a single group, the *Imitervirales*. We tested whether Larry and Curly followed this same pattern by infecting the freshwater alga *Chrysochromulina parva* with each PLV alone or together with one of two algal viruses known to infect *C. parva*. Neither PLV grew on its own, and neither influenced algal growth. They also failed to replicate when paired with the *Imitervirales* virus CpV-BQ3, showing that this known helper virus cannot support them. In contrast, both PLVs replicated when co-infecting with CpV-BQ1, a virus from the *Algavirales*. During this infection, CpV-BQ1’s own replication dropped, indicating a hyperparasitic interaction. These findings provide the first culture-based evidence that PLVs can use *Algavirales* viruses as helpers. They also show that *C. parva* hosts multiple, distinct layers of viral parasitism, expanding our understanding of the complexity and diversity of virus–virus interactions in microbial ecosystems.

## Introduction

Virophages were first reported in 2008 with the discovery of Sputnik alongside its mamavirus helper virus, which infects the protist *Acanthamoeba castellanii*. This discovery established the existence of a tripartite system consisting of a virophage, a giant helper virus, and a shared host (1). Since Sputnik’s discovery, additional tripartite systems involving satellite viruses with virophage-like replication have been identified in freshwater and marine environments worldwide (e.g., 2, 3-5). The term *virophage* has been used informally to describe dsDNA viruses that hyperparasitize helper dsDNA viruses of the phylum *Nucleocytoviricota* and formally refer to viruses classified within the class *Virophaviricetes* (phylum *Preplasmiviricota)* (6).

Related to virophages, and engaging in similar hyperparasitic replication strategies, are the polinton-like viruses (PLVs) which share ancestry with mobile genetic elements known as polintons. Unlike polintons, PLVs encode major capsid proteins (MCP) characteristic of dsDNA viruses in the phylum *Nucleocytoviricota* (7, 8). As with virophages in the *Virophaviricetes*, only a few PLVs have been isolated; most are known from genomic and metagenomic sequences (e.g., 9, 10-13).

Given the rapid pace of discovery of virophages, it is not surprising that the taxonomy of these hyperparasites has been revised in recent years (6, 7, 14). Despite taxonomic distinctions between PLVs and virophages (*sensu stricto*), cultured representatives of both groups are associated with helper viruses belonging to the *Imitervirales* families *Mesomimivirdae* and *Mimiviridae* (14-16). To date, no virophage or virophage-like virus has been experimentally shown to require a phycodnavirus for its replication. Genomic evidence from the Organic Lake Antarctica virophage (17) and tripartite systems identified in Dishui Lake, China (18) raised the possibility that some virophages parasitize members of the *Phycodnaviridae*, but these associations were inferred only from metagenomics. Additionally, some viruses that were loosely considered phycodnaviruses because they infect phototrophic protists were later classified in the *Imitervirales* (19, 20).

The freshwater prymnesiophyte *Chrysochromulina parva* is infected by viruses of the *Algavirales* and *Imitervirales*, and PLVs (21-23). Similarly, both *Algavirales* and *Imitervirales* viruses are known to infect the marine prymnesiophyte algae *Phaeocystis globosa* (PgVs), and the *Imitervirales* PgV-16T and PgV-14T (24, 25) are themselves parasitized by PLVs. These observations suggest that infections by a consortium of viruses is a common feature of haptophyte ecology that predates the transition of some haptophytes from marine to freshwater environments (26).

Studies of the *C. parva–*virus system demonstrated the virophage-like replication of a putative PLV, CpV-PLV Moe (22) provided evidence that Moe’s replication depends on CpV-BQ3, a strain of the *Imitervirales* species *Tethysvirus ontarioense*. CpV-PLV Moe behaves as a genuine virophage, inhibiting the replication of its helper CpV-BQ3 and supporting the growth of *C. parva* (22). Two additional PLVs, CpV-PLV Larry and Curly, were discovered during analysis of the *Tethysvirus ontarioense* genome (23). In experiments to isolate CpV-PLV Moe, cultures containing CpV-PLV Larry and Curly were generated alongside CpV-BQ1, the unclassified phycodnavirus originally described for *C. parva* (21, 27). This raised the question of whether these PLVs depend on a *Phycodnaviridae* helper rather than a *Mesomimivirdae* helper as is the case for Moe (22). In this study we infected *C. parva* with CpV-PLV Larry and Curly in the presence of either BQ1 or BQ3 and present the first culture-based evidence of a PLV parasitizing a phycodnavirus.

## Results and discussion

Hereafter, CpV-BQ1, CpV-BQ3, CpV-PLV Larry, CpV-PLV Curly, and CpV-PLV Moe will be referred to as BQ1, BQ3, Larry, Curly, and Moe, respectively.

Infections with Larry or Curly alone had no discernible effect on the growth of *C. parva* cultures over the 21-day sampling period (p > 0.05; Fig 1A), with all treatments reaching a peak abundance of approximately 3.5 × 10^6^ cells mL^−1^ by Day 14. Quantitative PCR (qPCR)-inferred abundances of both Larry and Curly declined by roughly an order of magnitude from their initial inoculum (∼8 × 10^5^ and 1 × 10^5^ copies mL^−1^, respectively) to ∼1 × 10^4^ copies mL^−1^ by Day 21 (Fig 1B). These findings demonstrated that neither Larry nor Curly alone can replicate within *C. parva* or effect its growth, consistent with the PLV Gezel-14T and the virophage Sputnik (1, 25).

**Fig 1.**
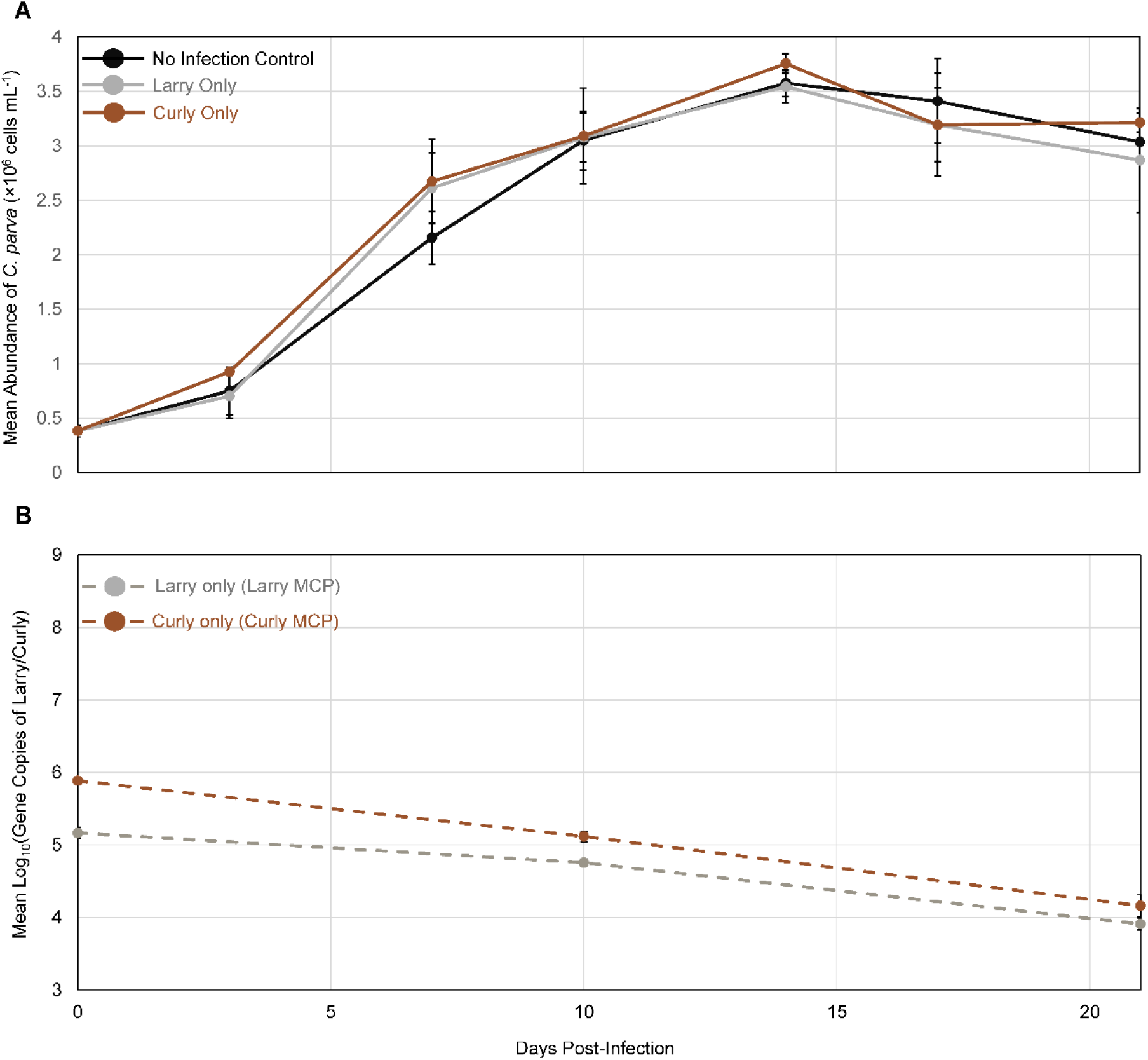
*C. parva* and virus abundances during incubations with each PLV. **(A)** Mean *C. parva* abundances (× 10^6^ cells mL^−1^) from triplicate flasks inoculated with Larry alone or Curly alone, alongside uninfected negative control flasks. **(B)** The abundances of Larry or Curly estimated by qPCR and shown as the mean log_10_ gene copies mL^−1^ of triplicate flasks. In both panels, the error bars represent the standard deviation of triplicate flasks, and where they are not visible, they were smaller than the data marker.

We next investigated BQ3, the helper virus of CpV-PLV Moe (22), as a candidate helper virus for Larry and Curly. All BQ3 treatments (alone or co-infected infecting with Larry or Curly) caused significant host lysis within 7 days, with cell counts dropping to ∼10^5^ cells mL^−1^ compared to ∼2 × 10^6^ cells mL^−1^ in no-infection controls (Fig 2A). Lysis coincided with a 3-order-of-magnitude increase in BQ3 abundance to ∼10^8^ copies mL^−1^ (Fig 2B). Neither Larry nor Curly affected BQ3 replication (p > 0.025; Fig 2B) or conferred protection against *C. parva* lysis (p > 0.05; Fig 2A). Both of these PLVs remained at initial inoculum levels (∼5 × 10^5^ to 10^6^ copies mL^−1^) throughout the 21-day incubation (p > 0.008; Fig 2B) indicating that the *Imitervirales* virus BQ3 is not their helper.

**Fig 2.**
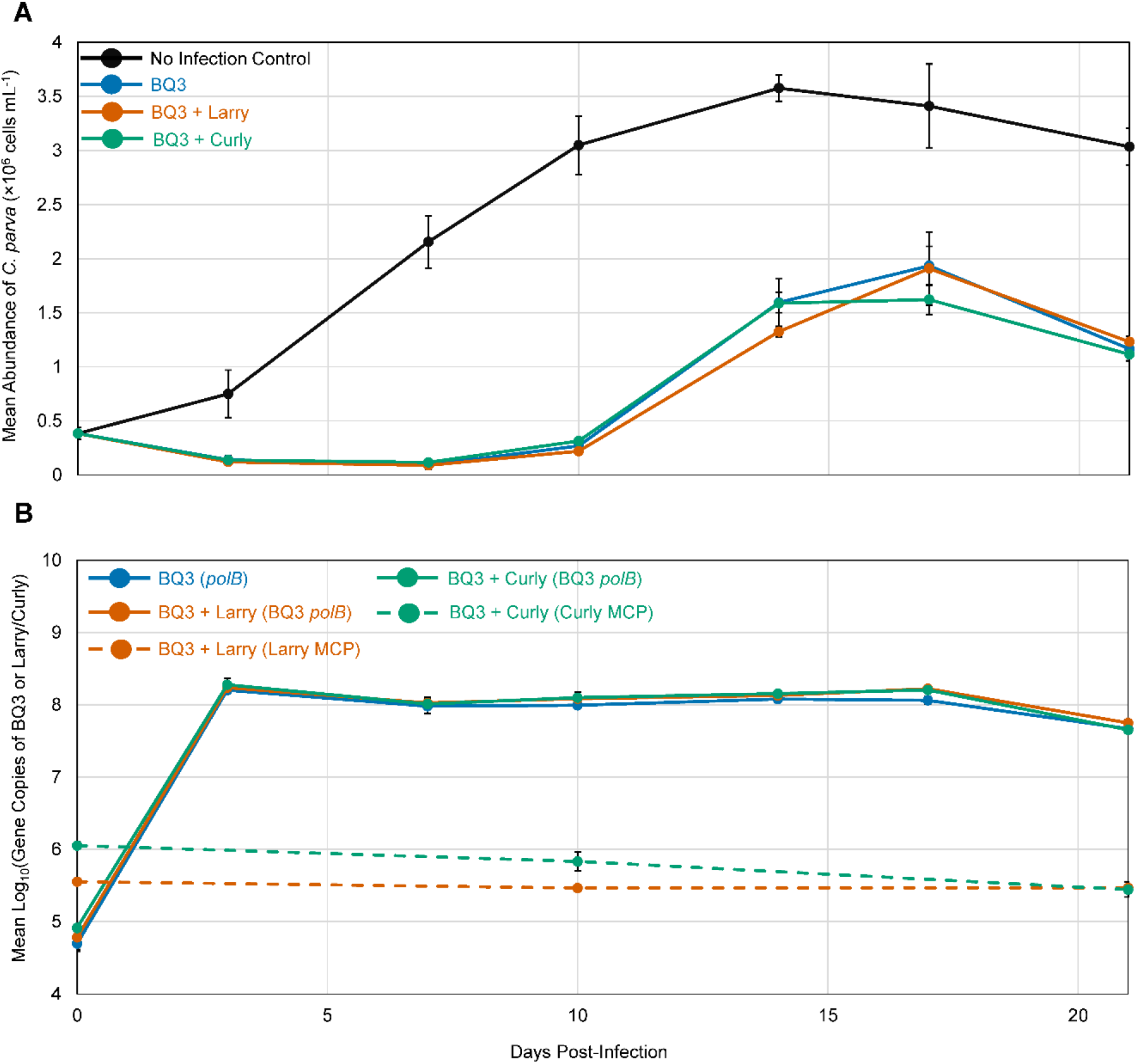
*C. parva* and virus abundances during infections with BQ3 and co-infections with Larry or Curly. **(A)** Mean *C. parva* abundance (× 10^6^ cells mL^−1^) from triplicate flasks co-inoculated with BQ3 and Larry or BQ3 and Curly, alongside flasks inoculated with BQ3 as an infection control as well as uninfected negative control flasks. **(B)** The abundance of BQ3 was estimated by qPCR targeting its *polB* gene (solid lines), whereas the abundances of Larry or Curly were estimated from qPCR of their major capsid protein (MCP) genes (dashed lines); values are mean log_10_ gene copies across triplicate flasks. For both panels, the error bars represent the standard deviation of triplicate flasks, and where they are not visible, they were smaller than the data marker.

We turned to BQ1, an unclassified phycodnavirus infecting *C. parva* (21, 23, 27), as a possible helper virus for Larry and Curly because it was co-purified alongside these PLVs. Using TEM, viral particles with expected sizes of ∼110 nm for BQ1 and ∼70 nm for Larry and Curly (16, 21, 22) were observed in these samples (Fig 3), which were also used as stocks for the mixed infections. Infections with BQ1 alone or co-infections with Larry or Curly inhibited *C. parva* growth, with an onset of lysis by 14 days post-infection (Fig 4A). After 21 days, the BQ1-only culture dropped to 9.5 × 10^5^ cells mL^−1^, compared to 3 × 10^6^ cells mL^−1^ in no-infection controls. Co-infection with Larry impeded cell lysis (1.5 × 10^6^ cells mL^−1^), whereas Curly slightly enhanced lysis (∼6.4 × 10^5^ cells mL^−1^); though neither difference was statistically significant (p > 0.05). Infections with BQ1 alone led to its 10,000-fold increase (∼10^5^ to ∼10^9^ copies mL^−1^) by day 21, whereas co-infection with either Larry or Curly limited BQ1 replication to a 100-fold increase (∼10^7^ copies mL^−1^; p < 0.008; Fig 4B). Notably, both PLVs replicated in the presence of BQ1, each increasing roughly 1000-fold from their initial inoculum (p < 0.025; Fig 4B).

**Fig 3.**
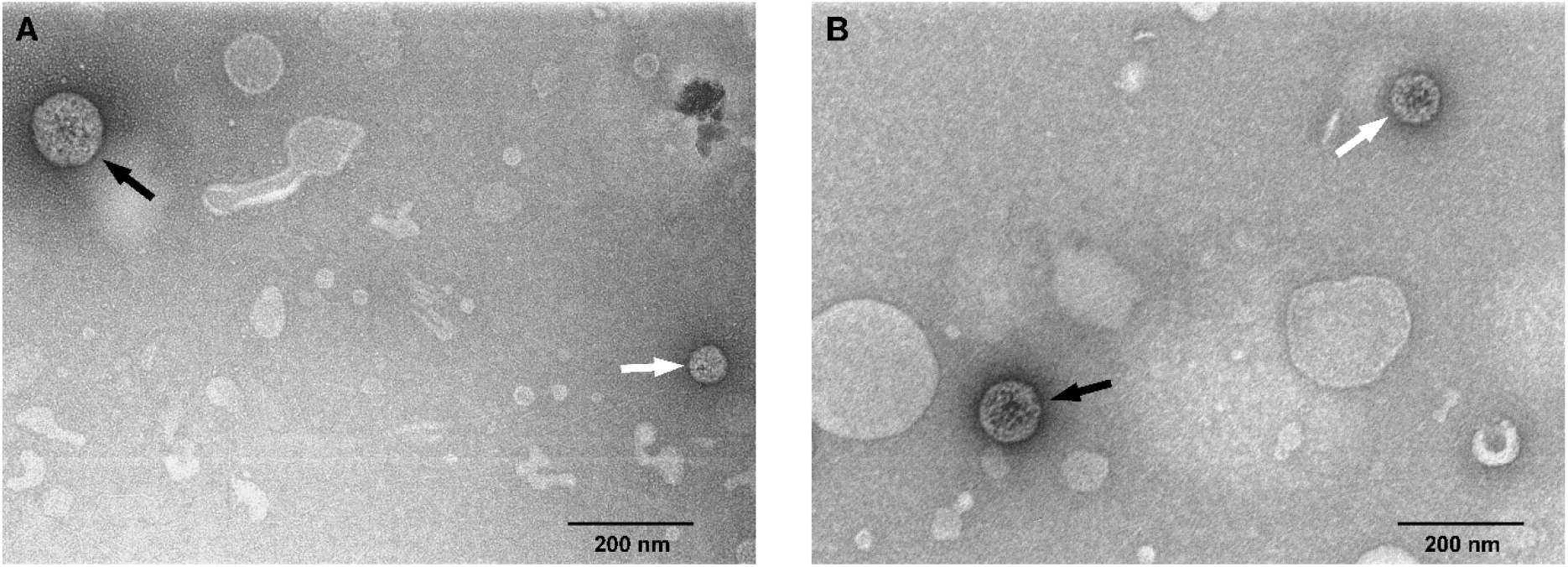
Transmission electron microscopy of viral lysates viewed at 30,000 X magnification. Samples applied to carbon-coated copper grids (FCF400-Cu-UB, Electron Microscopy Sciences, Hatfield, USA) were allowed to sit for 10 minutes, were washed with double-distilled H_2_O, and were stained using UranyLess EM Stain (Electron Microscopy Sciences, Hatfield, USA) for 1 minute before imaging. **(A)** Micrograph of the sample used for infection experiments with abundant CpV-BQ1 and CpV-PLV Curly as inferred from qPCR. The diameters of particles indicated with the solid black and solid white arrows are approximately 121 nm (range 118 to 123 nm depending on the axis measured) and 63 nm (range 61 to 65 nm), respectively. **(B)** Micrograph of the sample used for infection experiments with abundant CpV-BQ1 and CpV-PLV Larry. The diameters of the particles indicated with the solid black and solid white arrows are approximately 104 nm (range 102 to 106 nm) and 83 nm (range 82 to 84 nm), respectively.

**Fig 4.**
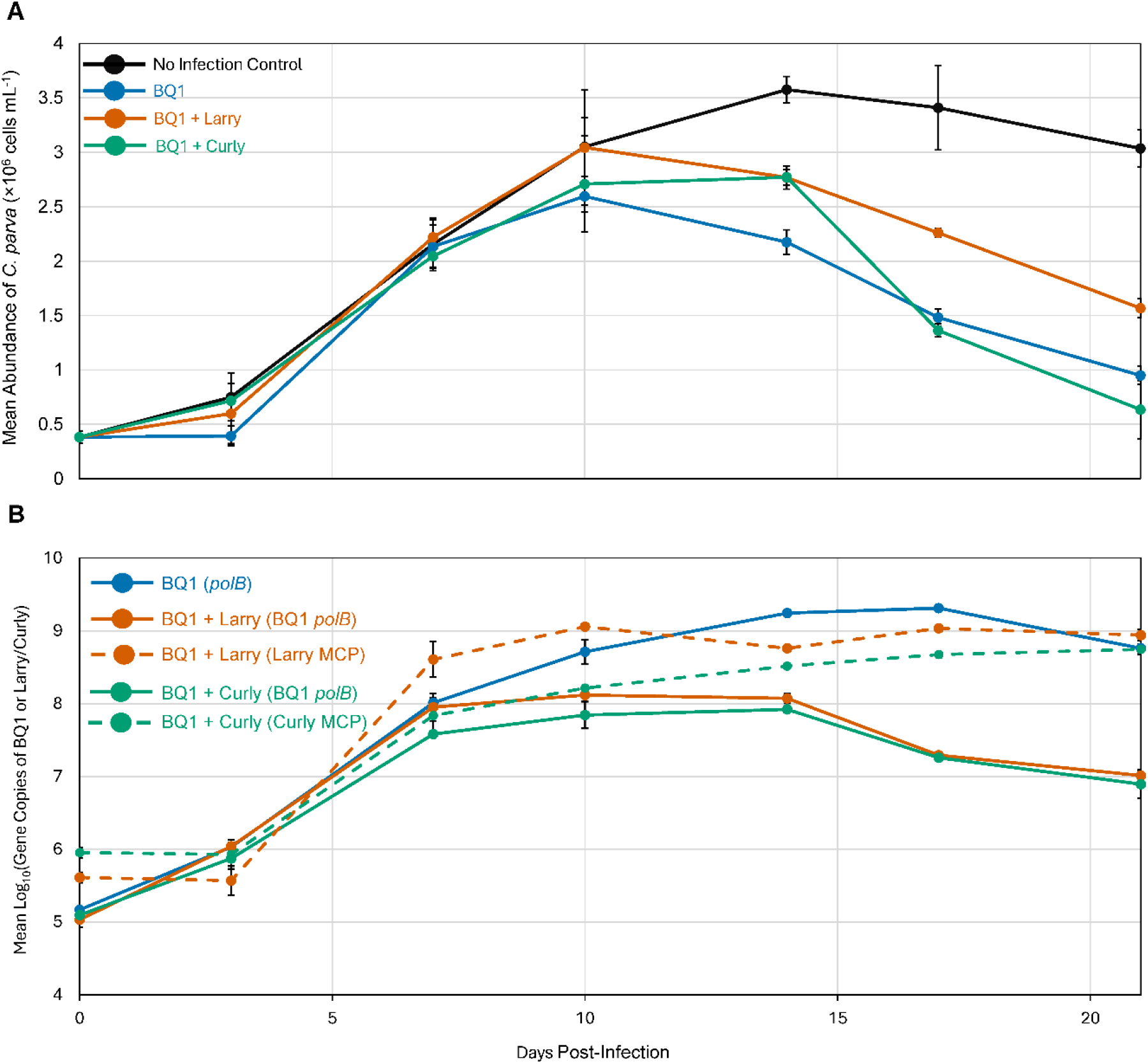
*C. parva* and virus abundances during infections with BQ1 and co-infections with Larry or Curly. **(A)** Mean *C. parva* abundance (× 10^6^ cells mL^−1^) in triplicate flasks co-inoculated with BQ1 and Larry or with BQ1 and Curly, alongside flasks inoculated with BQ3 as an infection control as well as uninfected negative control flasks. **(B)** The abundance of BQ1 was estimated by qPCR targeting its *polB* gene (solid lines), whereas the abundances of Larry or Curly were estimated from qPCR of their major capsid protein (MCP) genes (dashed lines); values are mean log_10_ gene copies across triplicate flasks. Error bars represent the standard deviation of triplicate flasks, and where they are not visible, they were smaller than the data marker.

Altogether, these results demonstrate that Larry and Curly are associated with a phycodnavirus, BQ1, as their helper virus. Their behavior is analogous to other known virophages associated with viruses in the *Imitervirales*. It appears that Larry may be able to mitigate BQ1-induced lysis, as seen in BQ3–Moe co-infections (22), whereas Curly does not, akin to Gezel-14T and its helper virus PgV-14T (25), though neither observation was statistically significant. Additional studies with varying densities of Larry or Curly relative to BQ1 could clarify the influence of these PLVs on their helper viruses and host cells.

This study provides the first evidence that the *C. parva*-virus system harbors at least two independent virophage–helper relationships spanning two distinct virus orders: *Algavirales* and *Imitervirales*. The observation that an isolated polinton-like virophage is both supported by and capable of inhibiting an *Algavirales* helper virus expands the known host range of virophages and polinton-like virophages beyond the *Imitervirales* (14-16).

## Materials and methods

### Cell and viral culture and quantification

Cell cultivation and infection experiments used non-axenic cultures of *Chrysochromulina parva* CCMP291 grown in DY-V medium (28) at 15 ^°C^ with a 12:12 h light/dark cycle at 23 μmol photons m^−2^ s^−1^ (22). Viral lysates were maintained by adding filtered lysates into a mid-log phase *C. parva* culture and, after cell lysis, filter-purifying the culture sequentially through 47 mm diameter, 0.50 μm nominal pore-size borosilicate glass microfiber filters (GC50, Advantec MFS, Inc. Dublin, USA), and 0.45 μm pore-size PVDF Durapore® membranes (MilliporeSigma, Burlington, USA). To further purify Larry and Curly from the lysates that also contained BQ1, whose particles are just above 0.1 μm in diameter (21), lysates were serially filtered six times through a 0.1 μm pore-size PVDF Minisart® syringe filter (Sartorius, Shinagawa City, Japan).

*C. parva* cell abundances and virus particle abundances were enumerated as previously described (22). Briefly, cells were counted using a haemocytometer and bright-field light microscope after staining with Lugol’s iodine. Virion abundance was inferred from gene copy abundances using the 5′ nuclease assay targeting PolB genes for BQ1 and BQ3, and the major capsid protein (MCP) genes for Larry and Curly. Quantitative PCR primers and probes, and PCR conditions are described in Thomas, Inamoto (22).

### Larry and Curly isolation and infection experiments

Previously described efforts to isolate BQ3 and Moe (22) also yielded a sample containing only BQ1, Larry, and Curly. Quantitative PCR showed Larry and Curly to be more abundant than BQ1 in this sample, hence each PLV was isolated by end-point dilution and was added to *C. parva* cultures inoculated with BQ1 as the presumptive helper. Samples confirmed by qPCR to contain only a single PLV were propagated for several generations using *C. parva* cultures infected with BQ1.

Experimental infections were set up in triplicate disposable 75 cm^2^, 250 mL capacity, vented, polystyrene cell culture flasks (Sarstedt, Montreal, Canada) at a starting *C. parva* abundance of ∼4.0 × 10^5^ cells mL^-1^. Trials included no-infection controls, infections with BQ1, BQ3, Larry, or Curly alone, and co-infections of BQ1 or BQ3 with either Larry or Curly. Viruses were added at approximately 1:1:1 ratio based on *C. parva* cell counts, viral *polB* copies, and PLV MCP gene copies. Culture flasks were arranged within the growth chamber to maximize variability of irradiance for each set of replicates. 200 μL samples were aseptically collected from each flask during sampling days; cell counts were performed immediately, whereas qPCR samples were stored at −20°C until further analysis.

### Electron Microscopy

The BQ1 and Larry or BQ1 and Curly samples used for infection experiments were concentrated by centrifugation, as described (22). Five μL of concentrated samples were applied to copper grids, which were then washed and stained before imaging with a HT7700 transmission electron microscope (Hitachi) at the Imaging Facility of the Department of Cell & Systems Biology, University of Toronto. Micrographs were analyzed with Fiji (29).

### Statistical Analyses

*C. parva* abundances were log10-transformed to improve normality and homoscedasticity, with residuals checked by Q–Q plots and Levene’s test. Abundances were compared by ANOVA with Tukey’s HSD at Day 21 for the negative control, Larry, Curly, BQ1, BQ1 + Larry, BQ1 + Curly treatments, and at Day 7 (following BQ3 lysis) for the negative control, BQ3, BQ3 + Larry and BQ3 + Curly treatments. Day 21 BQ1 and BQ3 abundances were each compared against their respective co-infection treatments (Bonferroni-adjusted α = 0.025). To assess Larry and Curly replication, log10-transformed Day 0 abundances were subtracted from Day 21 abundances for six treatments (Larry and Curly alone and each with BQ1 or BQ3), and one-sided, one-sample t-tests determined whether these changes exceeded zero (Bonferroni-adjusted α = 0.0083).

## Acknowledgments

The authors wish to acknowledge Dr. Kenana Al Kakouni, the Lead Microscopy Technician at the Imaging Facility of the Department of Cell and Systems Biology at the University of Toronto, for their assistance with transmission electron microscopy.

